# Expression and Purification of a Mammalian Protein: Cytosolic Domain of IRE1α from Insect Sf21 Cells

**DOI:** 10.1101/750430

**Authors:** Amrita Oak, Grace Jansen, Christina Chan

## Abstract

Eukaryotic proteins can be expressed in different heterologous systems. However, mammalian proteins in general have specific post-translational processing requirements that may not be fulfilled by a regular bacterial expression system. In this study, we use an insect cell system to express a mammalian protein of interest. Spodoptera frugiperda (Sf21) cells were used in conjunction with a baculoviral expression system to produce the cytosolic domain (CD) of IRE1, an endoplasmic reticulum (ER) stress sensor protein. Inositol Requiring Enzyme 1 (IRE1) is a dual function kinase and endoribonuclease protein that cleaves X-box binding protein (XBP1) mRNA. We used the pFastBac plasmid to insert the coding sequence into a recombinant bacmid shuttle vector which was then used to infect Sf21 cells. The expressed protein was then purified with an MBPTrap column to obtain >85% pure protein.

## 1. Introduction

Heterologous expression of proteins is defined as the expression of proteins in an organism that does not naturally express the protein. The different systems in use for such purposes can be broadly classified as bacterial, yeast, insect and mammalian systems. Cells in culture can be manipulated to express the protein of interest in transient or stable mode. Some factors that need to be considered while choosing the expression system are the required yield of the protein, post-translational modifications, functionality, and speed of expression.

Proteins of prokaryotic origin are best expressed in bacterial systems. *E. coli* can express and adequately process prokaryotic proteins and is the go-to system for cheap, scalable and high yield expression. If the expression of eukaryotic proteins is required, additional factors need to be taken into consideration. The most important factor would arguably be the level of post-translational modifications (PTM) required for a functional form of the protein. Proteins requiring extensive PTMs will not be processed correctly in bacterial systems and will most likely aggregate in inclusion bodies[1]–[3]. Adding a fusion protein tag to the protein of interest may sometimes help to resolubilize the proteins. However, in the interest of saving time and effort, it is advisable to switch to higher eukaryotic systems such as insect or mammalian cells. Another advantage to eukaryotic systems is the presence of extensive protein folding machinery that is vital for the function of the protein [4].

**Table 1** compares the different heterologous protein expression systems with respect to the type of protein, post-translational modifications and ease of large-scale production [5]– [10].

**Table 1:**
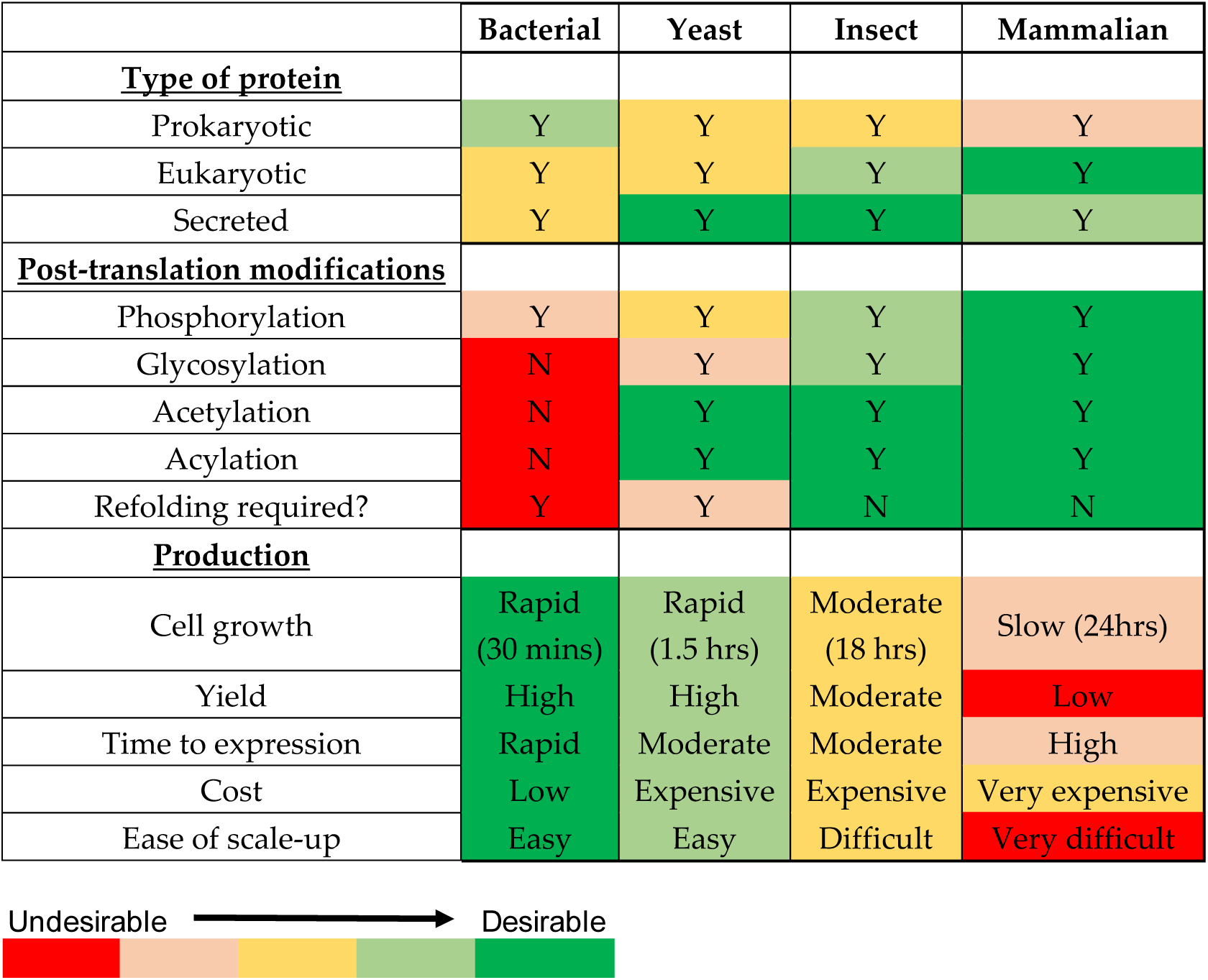
Characteristics of heterologous protein expression in bacterial, yeast, insect and mammalian cell systems ranked according to desirability

Bacterial systems have desirable characteristics for large scale production but are not able to process and correctly fold proteins requiring extensive PTMs [11]. Depending on the application of the protein, Chinese hamster ovary (CHO) or Human embryonic kidney (HEK 293) cells are typically the system of choice for eukaryotic proteins [12]. This is true, especially for therapeutic proteins or antibodies. However, the yield is very low compared to insect or bacterial cells – typically on the order of 100mg/L of expressed protein [13]. The level of protein expression is higher in insect cells up to 100mg/ L [14], [15].

Insect cells and mammalian systems have a wide range of post-translational modifications. An excellent tool to determine all possible post-translational modifications of a protein of interest is dbPTM (http://dbptm.mbc.nctu.edu.tw/). Insect and mammalian systems are both capable of all PTMs with one exception. Insect cells and mammalian cells have similar phosphorylation and O-linked glycosylation patterns. They can authentically process phosphorylation modifications, partly due to the presence of phosphatases. However, in terms of N-glycosylation, proteins in insect cells are high-mannosylated while mammalian cells have a complex glycosylation pattern [4], [12]. **Figure 1** depicts the difference in N-linked glycosylation patterns in insect cells vs. mammalian cells. Regular insect Sf9 and Sf21 cells will have a high mannose pattern of glycosylation but mimic Sf9 (available from ThermoFisher) cells are engineered to make complex N-glycans with terminal sialic acid. Varied and complex sugars such as N-acetylneuraminic acid, galactose, fucose, mannose are added in mammalian cells in a branched configuration as opposed to the addition of only mannose residues in insect cells.

**Figure 1:**
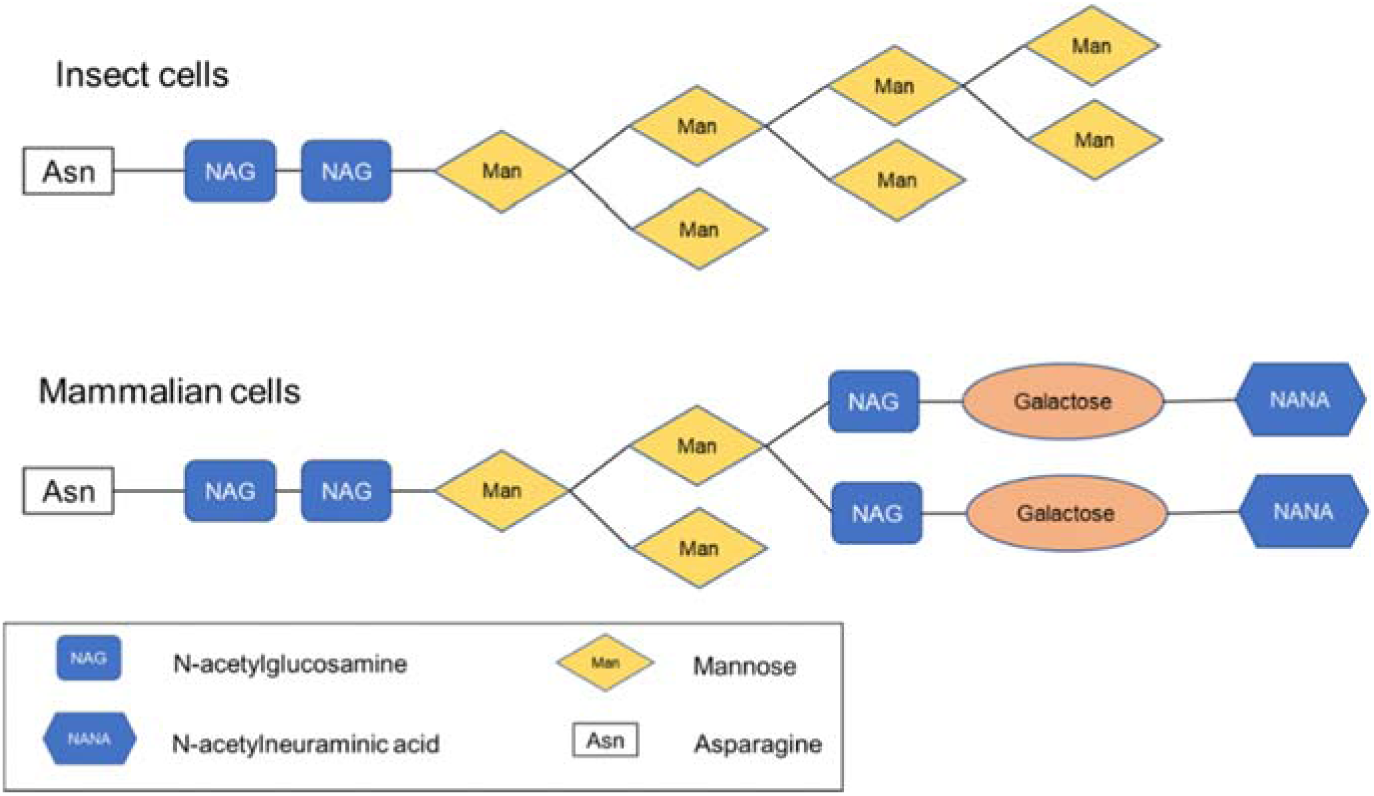
N-linked glycosylation patterns for insect cells and mammalian cells.

In this study, we developed a protocol to express the Inositol Requiring Enzyme-1α (IRE1α) protein, an ER stress sensor. IRE1 is a type I transmembrane protein primarily composed of three domains, the luminal domain, a transmembrane domain and cytosolic domain (CD)[16].

IRE1α is a bifunctional protein with kinase and endoribonuclease activity on the CD. We have developed a protocol to produce IRE1α-CD (547-977 aa) in insect Sf21 cells using baculovirus expression. IRE1α has been expressed in *E. coli* cells in our lab (unpublished data) as well as by other groups[17], [18]. However, IRE1α expressed in *E. coli* and yeast systems is in a hyperphosphorylated state[18], [19]. All the purification methods used thus far for purifying this protein have used the 6X His tag. The cell line used was Sf9 [18], [22]. However, there are no detailed methods or protocol papers published for this procedure, that is, using the Sf9 cells and 6X His tag method.

Post-translational modifications for IRE1 were predicted using algorithms developed by the Center for Biological Sequence Analysis, Technical University of Denmark (http://www.cbs.dtu.dk/ [1]. The predicted N-glycosylation, O-glycosylation, C-mannosylation and glycation sites are scored for the likelihood of modification (**Table 2**). The scores that cross the threshold (which is set by the algorithm at 0.5) are denoted as “positive” hits.

**Table 2:** Predictions for O-glycosylation, C-mannosylation, N-glycosylation and glycation sites on CD-IRE1.

| O-glycosylation |  |  |  | C-mannosylation |  |  |  |
| --- | --- | --- | --- | --- | --- | --- | --- |
| start | end | score | comment | start | end | score | comment |
| 2 | 2 | 0.792 | #POSITIVE | 191 | 191 | 0.18 | . |
| 3 | 3 | 0.796 | #POSITIVE | 287 | 287 | 0.271 | . |
| 5 | 5 | 0.791 | #POSITIVE |  |  |  |  |
| 15 | 15 | 0.189 |  |  |  |  |  |
| 16 | 16 | 0.506 | #POSITIVE |  |  |  |  |
| 24 | 24 | 0.111 |  |  |  |  |  |
| 38 | 38 | 0.015 |  |  |  |  |  |
| 61 | 61 | 0.043 |  |  |  |  |  |
| 73 | 73 | 0.040 |  |  |  |  |  |
| 85 | 85 | 0.006 |  |  |  |  |  |
| 102 | 102 | 0.018 |  |  |  |  |  |
| 121 | 121 | 0.110 |  |  |  |  |  |
| 126 | 126 | 0.102 |  |  |  |  |  |
| 127 | 127 | 0.213 |  |  |  |  |  |
| 128 | 128 | 0.181 |  |  |  |  |  |
| 135 | 135 | 0.052 |  |  |  |  |  |
| 151 | 151 | 0.107 |  |  |  |  |  |
| 164 | 164 | 0.034 |  |  |  |  |  |
| 178 | 178 | 0.107 |  |  |  |  |  |
| 180 | 180 | 0.741 | #POSITIVE |  |  |  |  |
| 183 | 183 | 0.643 | #POSITIVE |  |  |  |  |
| 188 | 188 | 0.111 |  |  |  |  |  |
| 198 | 198 | 0.148 |  |  |  |  |  |
| 206 | 206 | 0.026 |  |  |  |  |  |
| 208 | 208 | 0.047 |  |  |  |  |  |
| 213 | 213 | 0.013 |  |  |  |  |  |
| 223 | 223 | 0.070 |  |  |  |  |  |
| 226 | 226 | 0.040 |  |  |  |  |  |
| 232 | 232 | 0.064 |  |  |  |  |  |
| 245 | 245 | 0.252 |  |  |  |  |  |
| 276 | 276 | 0.510 | #POSITIVE |  |  |  |  |
| 288 | 288 | 0.454 |  |  |  |  |  |
| 300 | 300 | 0.102 |  |  |  |  |  |
| 300 | 300 | 0.102 |  |  |  |  |  |

| Glycation |  |  |  |
| --- | --- | --- | --- |
| start | end | score | comment |
| 22 | 22 | -0.943 | . |
| 28 | 28 | -0.707 | . |
| 53 | 53 | 0.801 | YES |
| 87 | 87 | -0.875 | . |
| 110 | 110 | 0.832 | YES |
| 144 | 144 | -0.529 | . |
| 158 | 158 | 0.699 | YES |
| 160 | 160 | 0.863 | YES |
| 170 | 170 | -0.773 | . |
| 171 | 171 | 0.83 | YES |
| 202 | 202 | -0.95 | . |
| 231 | 231 | -0.962 | . |
| 253 | 253 | -0.799 | . |
| 265 | 265 | 0.806 | YES |
| 273 | 273 | -0.94 | . |
| 282 | 282 | -0.878 | . |
| 291 | 291 | -0.83 | . |

| N-glycosylation |  |  |  |
| --- | --- | --- | --- |
| start | end | score | comment |
| 204 | 204 | 0.5427 | + |

## 2. Experimental Design

### 2.1 Materials

All the materials are listed separated by each sub-section of the procedure.

#### Cloning of IRE1α-CD into pFastBac plasmid

i. pFastBac His6 MBP N10 TEV LIC cloning vector (4C) (Addgene plasmid #30116)
ii. SspI-HF (New England Biolabs, cat # R3132S)
iii. CutSmart buffer (New England Biolabs cat# B7204S)
iv. QIAquick PCR Purification Kit (Qiagen, cat# 28104)
v. Deoxynucleotide (dNTP) set includes dGTP, dCTP (New England Biolabs, cat # N0446S)
vi. Bovine Serum Albumin (BSA), Molecular Grade (New England Biolabs, cat# B9000S)
vii. T4 DNA Polymerase (New England Biolabs, cat# M0203S)
viii. Q5 High Fidelity 2X Master Mix (New England Biolabs, cat# M0492S)
ix. OneShot Top10 chemically competent *E. coli* DH5α (ThermoFisher, cat# C404003)
x. Luria Broth (Sigma-Aldrich, cat# L3397)
xi. Luria Agar (Sigma-Aldrich, cat# L3272)
xii. LB agar plates containing ampicillin (Sigma-Aldrich, cat # L5667-10EA)

#### Preparation of recombinant bacmid

xiii. MAX Efficiency *E. coli* DH10Bac competent cells (ThermoFisher cat# 10361012)
xiv. Antibiotics and their stock solutions are listed in **Table 3**. Aliquot and store at −20°C. **Table 3:**
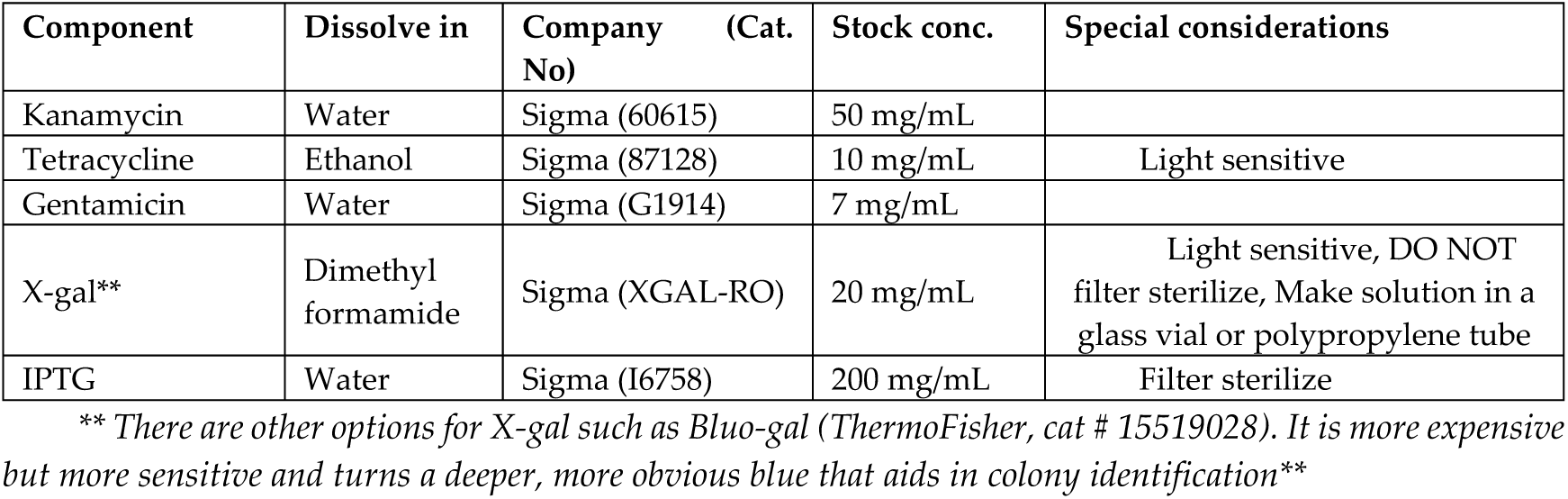
Antibiotic concentrations and stock solutions.
xv. S.O.C medium (ThermoFisher cat# 15544034)
xvi. PureLink HiPure Plasmid Miniprep Kit (Invitrogen, cat# K210002)

#### Transfection of recombinant bacmid into Sf21 cells

i. Sf-900-III serum-free media (ThermoFisher, cat# 12658019)
ii. Cellfectin II reagent (ThermoFisher, cat# 10362100)
iii. Gelcode Blue Stain Reagent (ThermoFisher, cat# 24590)
iv. IRE1α (14C10) Rabbit mAb (Cell Signaling Tech, cat# 3294)

#### Protein purification using MBPTrap column

v. Xtractor cell lysis buffer (Takara, cat# 635671)
vi. cOmplete, Mini, EDTA-free protease inhibitor cocktail (Sigma, cat# 11836170001)
vii. MBPTrap HP column (GE, cat# 28-9187-78)
viii. D-(+)-Maltose monohydrate (Sigma, cat# 63418-25G)
ix. Tris-Cl (Sigma, cat # 10812846001)
x. NaCl (Sigma, cat # S3014)
xi. Dithiothreitol (DTT) (Sigma, cat# DTT-RO)
xii. EDTA (Sigma, cat # 324504)
xiii. Phosphate buffered saline (PBS) (Sigma, cat# P7059-1L)
xiv. Binding buffer for MBPTrap HP column: 20 mM Tris-HCl, 200 mM NaCl, 1 mM DTT, 1 mM EDTA, pH 7.4
xv. Elution Buffer for MBPTrap HP column: Binding buffer + 10mM maltose
xvi. Regeneration buffer: 0.5M NaOH

## 3. Procedure

**Figure 2** depicts a flowchart of the steps involved in the expression and purification of IRE1α-CD in Sf21 cells using baculoviral vectors. The procedure is divided into 5 subunits. The time required for each subunit is indicated in brackets in the left column (**Figure 2**).

### 3.1 Cloning of IRE1α-CD into pFastBac plasmid

#### Note

pFastBac His6 MBP N10 TEV LIC cloning vector (4C) was a gift from Scott Gradia (Addgene plasmid #30116; http://n2t.net/addgene:30116; RRID: Addgene_30116). Another version of this plasmid (5C) is available so that two proteins can be expressed simultaneously. The sequence for IRE1α-CD was obtained from NCBI (Gene id: 2081). Primers were designed to clone the CD fragment from 547 aa-977 aa from a pCDNA-hIRE1 plasmid containing a full-length IRE1 coding sequence.

**Figure 2:**
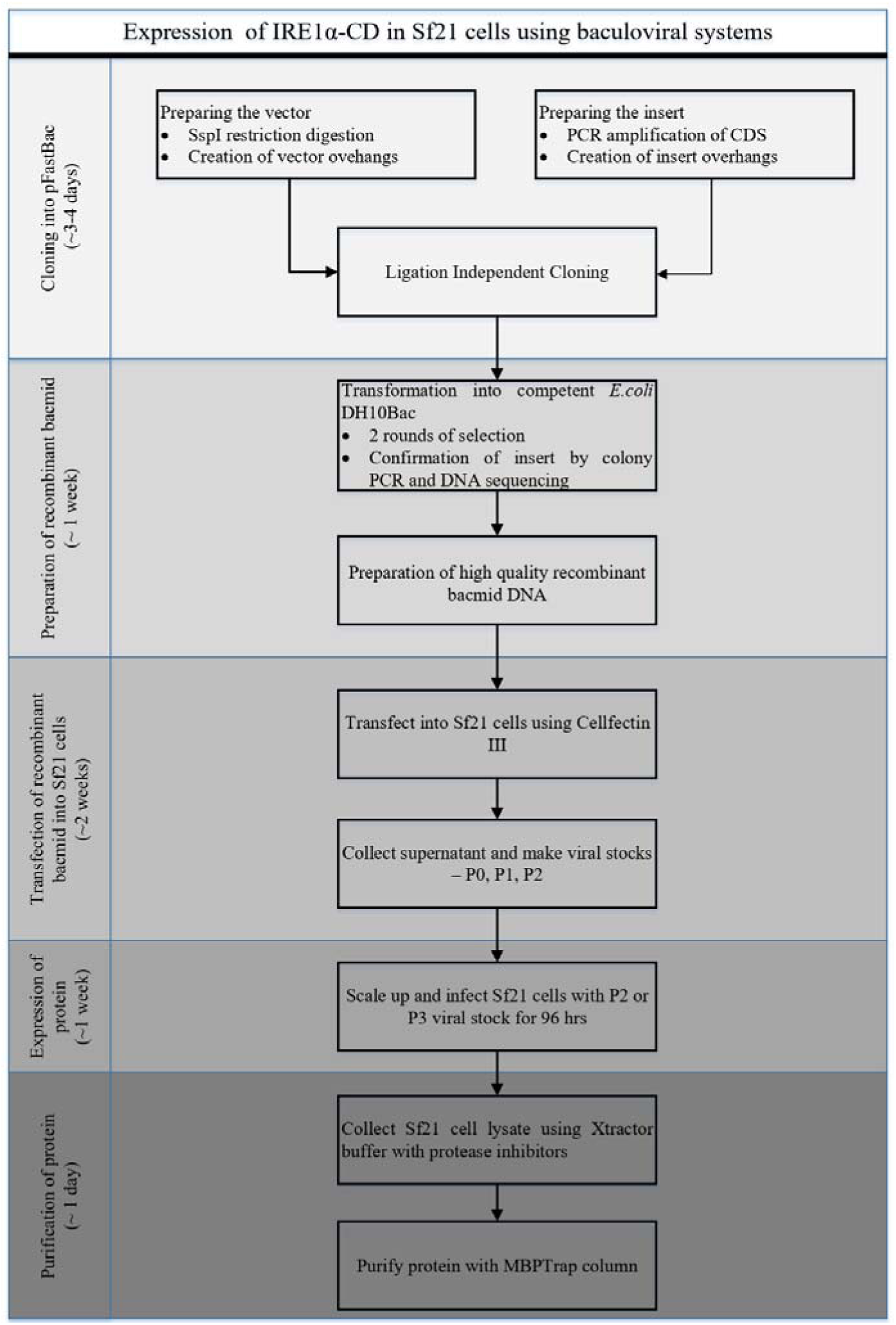
Flowchart for all the steps involved in the expression and purification of proteins from Sf21 cells.

**Figure 3:**
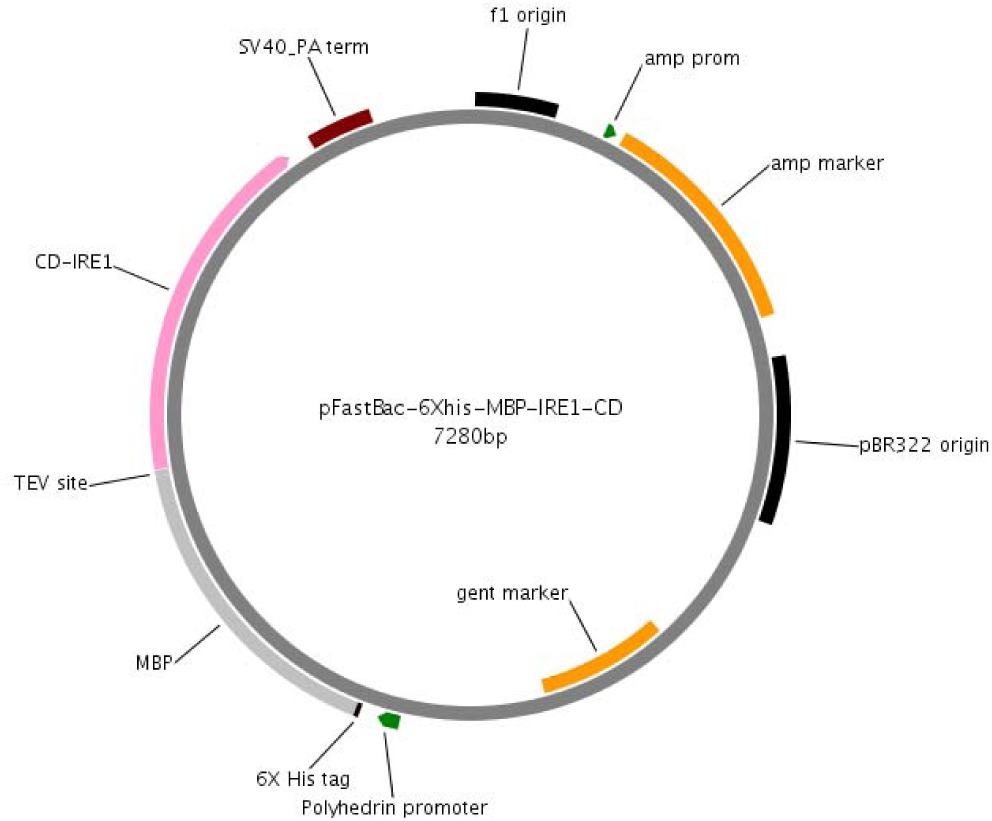
Plasmid map for pFastBac-IRE1α-CD.

**Table 4** shows the primers designed for the insertion of IRE1α-CD into the empty pFastBac plasmid. The underlined tag on each primer is to facilitate ligation independent cloning (LIC) and is specific to the pFastBac backbone. The rest of the primer is specific to the insert coding sequence (CDS). The LIC technique makes use of the endonuclease activity of T4 DNA polymerase to generate sticky overhangs for ligation between the vector plasmid and insert DNA. This technique avoids the use of ligases.

##### Protocol

###### Preparing the vector pFastBac plasmid

i. Linearize the plasmid with SspI-HF (NEB) (**Table 5**). The high-fidelity (HF) versions of restriction enzymes are faster and more efficient. Incubate at 37°C for 15 mins – 2 hrs to completely digest all the plasmid DNA. Heat inactivation is performed for 20 mins at 65°C.
ii. Use a PCR purification kit (QIAquick) to purify the large linearized fragment of plasmid DNA.
iii. To create pFastBac vector overhangs, mix the following components in PCR tubes, the order of addition is as listed (T4 DNA polymerase is added last) (**Table 6**). Incubate at 12°C for 30 minutes followed by heat inactivation at 75°C for 20 minutes.

**Table 4:**
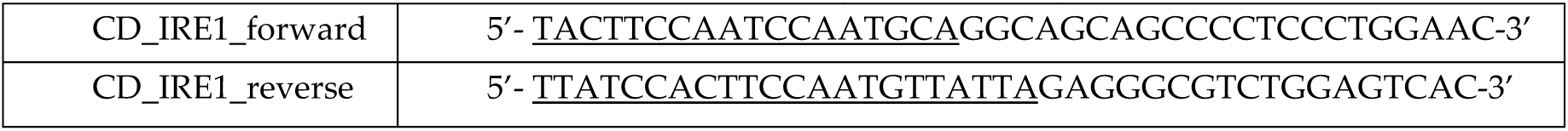
PCR primers for insertion of IRE1-CD (547aa-977aa) into pFastBac plasmid.

**Table 5:** Restriction digestion with SspI-HF.

| Component | Amount | Volume (μL) |
| --- | --- | --- |
| pFastBac plasmid DNA (250ng/μL) | 1 μg | 4 |
| 10X CutSmart buffer | Final concentration 1X | 5 |
| SspI-HF | 10 units | 1 |
| Water | Make up to 50 μL | 40 |

**Table 6:** Mix to create pFastBac vector overhangs.

| Reagent | Final conc. | Volume (μL) |
| --- | --- | --- |
| 10x NEB 2.1 | 1x | 4 |
| Eluted linearized pFastBac DNA | 10-50 ng/ μL | 20-30 |
| dGTP (100mM) | 2.5 mM | 1 |
| DTT (100mM) | 5 mM | 2 |
| BSA (10 ug/μL) | 0.25 μg/ μL | 0.6 |
| T4 DNA polymerase | 0.075 units/ μL | 1 |
| Sterile dH<sub>2</sub>O to 40ul |  |  |

###### Preparing the insert IRE1α-CD

iv. Set up a PCR to amplify the IRE1α-CD insert using the primers in **Table 4** as outlined in **Table 7**. Confirm amplification by running the PCR product on a 1.5% agarose gel. **Table 7:**
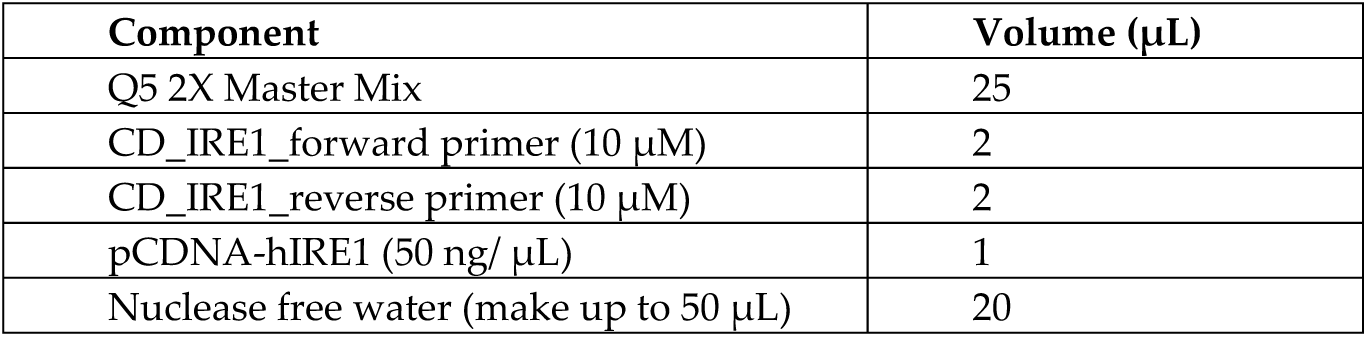
PCR mix for preparing IRE1α-CD insert. The PCR program should be run for 98°C 1 min, followed by 35 cycles of 98°C for 30 sec, 61°C for 10 sec, 72°C for 90 sec, and a final extension of 72°C for 5 mins.
v. Create the insert overhangs in a way similar to the vector overhangs in the presence of dCTP (instead of dGTP).
vi. After heat inactivation, mix both vector and insert together (1:3 ratio). The total volume of the combined vector and insert should be between 5-10 uL. Incubate for 5 mins at room temperature and then add 1uL of 25 mM EDTA followed by another incubation for 5 mins.
vii. Transform 2 µL of this mixture into competent OneShot Top10 *E. coli* DH5α (100 µL), spread onto LB agar plates containing 100 µg/mL ampicillin and incubated at 37°C for 16-24 hrs.
viii. Select individual colonies (5-10) and prep the plasmid DNA. Confirm the insertion of IRE1α-CD by DNA sequencing using primers specific to the IRE1α-CD sequence (seq for-5’-aagcagctccagttcttccaggac-3’)

### 3.2 Preparation of recombinant bacmid

#### Note

Once the insert has been cloned into the pFastBac empty plasmid, it needs to be transformed into *E. coli* DH10Bac cells to form the recombinant bacmid. *E. coli* DH10Bac competent cells are sold by ThermoFisher. The genotype is F-mcrA Δ (mrr-hsdRMS-mcrBC) F80lacZΔM15 Δ lac X74 recA1endA1 araD139 Δ (ara, leu)7697 galU galK λ - rps L nupG /pMON14272 / pMON7124. DH10Bac cells have a baculovirus shuttle vector and a helper plasmid. This machinery is required for the generation of recombinant bacmid after transformation of pFastBac-IRE1α-CD into the cells. The baculovirus shuttle vector (bMON14272) also encodes kanamycin resistance, and the helper plasmid (pMON7142) has tetracycline resistance. The presence of the Φ80lacZΔM15 Δ lac marker enables the use of blue/white colony screening to determine integration of the insert into the bacmid. pFastBac has the Tn7 element which includes the polyhedrin promoter, the gene of interest and gentamicin resistance.

It is also possible to use regular *E. coli* DH10Bac cells and make them chemically competent (see **Supplemental file**).

## Protocol

i. Dissolve LB agar powder in water and autoclave at 121°C for 15 mins.
ii. Let it cool to 55°C (there should be no lumps of agar).
iii. Add all the antibiotics and chemicals in the working concentrations listed in **Table 8**.
iv. Let harden, store at 4°C in the dark.
v. Thaw a vial of competent *E. coli* DH10Bac cells on ice in a 15 mL round bottom sterile polypropylene tube. \**Do not use 1.7 mL or 2 mL microcentrifuge tube*\*
vi. Add 100ng pFastBac-IRE1-CD plasmid DNA to the vial.
vii. Incubate on ice for **30** mins.
viii. Heat shock for **45 s** at **42°C**.
ix. Transfer tubes back on ice and chill for 5 mins.
x. Add **900 µL** SOC medium.
xi. Incubate tubes in a shaking incubator, 37°C at 200 rpm for at least **4 hrs**. \**This 4 hr long outgrowth step is necessary to allow the bacteria to generate the antibiotic resistance proteins encoded on the plasmid backbone*\*
xii. Make three 10-fold serial dilutions of 800 µL cells. Plate 100uL of each dilution onto LB plates with kanamycin, tetracycline, gentamicin, X-gal and IPTG.
xiii. Incubate plates at 37°C for **48 hrs** to allow enough time for colony formation and color development. Pick white colonies. \**Do not pick colonies before 48 hrs because it may be difficult to distinguish between white and blue colonies*\* \**Pick colonies that are large and well isolated. Avoid picking colonies that are gray or darker in the center as they may contain a mixture of empty bacmid and recombinant bacmid*\*
xiv. Replate selected colonies for an additional round of selection on fresh LB agar plates overnight at 37°C.
xv. Once the white phenotype is confirmed, inoculate in LB media with kanamycin, gentamicin, and tetracycline and grow overnight.
xvi. Isolate recombinant bacmid DNA using Purelink HiPure Plasmid Kit by Invitrogen with a modified protocol (see **Supplemental file**).

**Table 8:** Working concentrations of antibiotics.

| Component | Stock conc. | Working conc. | Dilution |
| --- | --- | --- | --- |
| Kanamycin | 50 mg/mL | 50 μg/mL | 1:1000 |
| Tetracycline | 10 mg/mL | 10 μg/mL | 1:1000 |
| Gentamicin | 7 mg/mL | 7 µg/mL | 1:1000 |
| X-gal | 20 mg/mL | 200 µg/mL | 1:100 |
| IPTG | 200 mg/mL | 40 µg/mL | 1:200 |

### 3.3 Transfection of recombinant bacmid into Sf21 cells

#### Note

Sf21 cells are suspension cells grown at 28°C, without the need for CO2 incubators. They are grown in Sf-900 III media. The media can be supplemented, if needed, with serum but bovine serum should not be used (fetal bovine serum (FBS) is preferred). The serum needs to be inactivated to inactivate complement fragments that can inactivate baculoviruses. Transfection reagents can be lipid based (e.g. cellfectin, fugene). Calcium chloride can be used for baculovirus transfection, but it has less efficiency than lipid-based reagents.

##### Protocol

###### Transfection of bacmid into Sf21 cells

i. A day before transfection, passage Sf21 cells so that they are at a density of 3 × 10^5^ cells/mL. \**Actively dividing cells have a higher transfection efficiency and produce more protein*\*
ii. Plate cells at a concentration of 8 x10^5^ cells/well of a 24-well plate.
iii. Allow the cells to attach for 1 hr and replace media with fresh media (without FBS).
iv. Use 2 µg of bacmid DNA per well. Dilute in 75 µL of Sf 900-III media. Incubate for 15 mins.
v. Use 8 µL of cellfectin II in 75 µL media. Incubate for 15 mins. \**Make sure the cellfectin II is mixed thoroughly before use, but do not vortex the tube*\*
vi. Mix together the two tubes and incubate for 15-30 mins.
vii. Add mixture dropwise to the well<u>s</u> of the 24-well plate.
viii. 24 hrs later, replace transfection media with fresh SF-900 III media containing 10% FBS. \**Serum proteins in FBS act as substrates for proteases*\*
ix. Check for signs of infection (SIF) after 24 hrs up to 96 hrs. \**SIF may not be obvious in P0 infection so continue infection for 5 days and collect the supernatant and re-infect for 5 more days to amplify the viral stock*\*
x. After changing media, collect supernatant after every 24 hrs for 24, 48, 72 and 96 hrs. This supernatant is the P0 viral stock. Store at 4°C in the dark.
xi. Use the P0 viral stock to infect newly plated Sf21 cells to generate P1 stock. Add 150 µL of P0 stock dropwise on top of the cells (plated in a 6-well plate), gently swirl a few times and incubate the plate at 28°C.
xii. Look for SIF in 24 hours post infection time. After 5 days, collect only P1 viruses with SIF and store viral stocks in the dark at 4°C.
xiii. In a similar fashion, collect P2, P3 and P4 stocks for increased baculoviral titer with an increasing number of cells infected (6-well plate → 150 cm dish→ T25 flask). Store all at 4°C protected from light.

###### Scale-up of heterologous protein expression

i. Seed Sf21 cells in a T75 (vented) flask as described in the transfection procedure. Make sure they are actively dividing cells in the log phase of growth.
ii. Add 150 µL of the collected P1 stock.
iii. Incubate at 27°C for 5 days or until 30-40 % of the cells have lysed.
iv. Collect cells as well as supernatant. **This will be the P2 viral stock**.
v. Use P2 viral stock to infect spinner flasks with Sf21 cells.
vi. If no signs of infection are observed, infect Sf21 cells with the P2 stock of baculoviruses and collect the P3 viral stock (see **Figure 5** for SIF). We concentrated the baculovirus viral supernatants with successive passages (P1,P2,P3) and used the passage that gave us visible signs of infection at 24 hrs with expression of protein. The viral titer (expressed as plaque-forming units/mL or pfu/mL) can also be determined with a plaque assay. Briefly, cells in a tissue culture dish are infected and overlayed with agarose. After the cells are grown for about 10 days, the plaques can be counted to determine the pfu/mL concentration.

**Figure 4:**
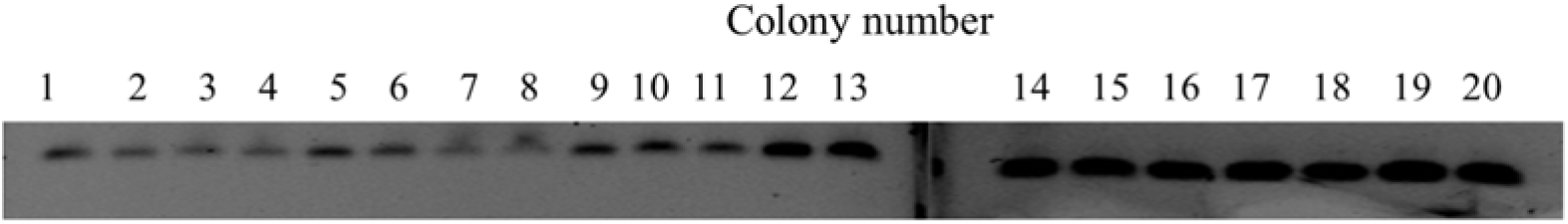
Colony PCR of white *E. coli* transformant colonies

**Figure 5:**
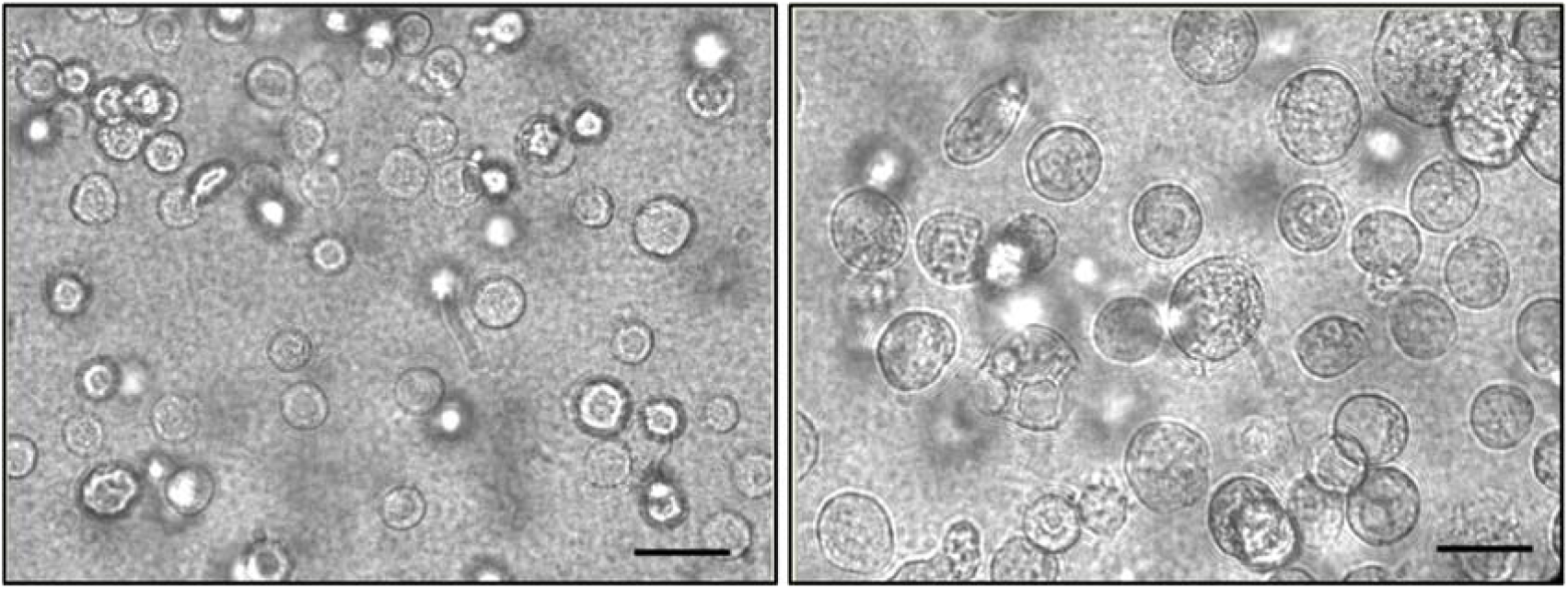
Signs of infection in untransfected Sf21 cells (left) and P3 treated Sf21 cells (right) for 24 hrs. Scale bar = 45µm

### 3.4 Purification of MBP fusion protein

#### Note

For this protocol, use Sf21 cells infected with P3 stock of baculovirus for 96 hrs. The cell pellet is collected and lysed with Xtractor buffer, and an MBPTrap column is used to extract MBP-tagged IRE1α-CD. The MBP tag can be cleaved from the protein by incubating with 1 unit of AcTEV protease (ThermoFisher, cat# 12575015) for 1 hr at 30°C for each 3µg protein.

##### Protocol

i. Prepare Xtractor buffer with protease inhibitor of choice. In this purification protocol, we used cOmplete EDTA free protease inhibitor cocktail.
ii. Pellet Sf21 cells at 100 × g for 15 mins, remove supernatant media and wash pellet once with phosphate buffered saline (PBS). Add 5 mL Xtractor buffer to 1 g cell pellet.
iii. Mix vigorously by vortexing and rock at room temperature for 15-30 mins. Make sure that cell lysis is complete by observing under the microscope.
iv. Centrifuge at 16,000 g for 20-30 mins to pellet the cell debris.
v. Take the clarified supernatant and filter through a 0.22 µm filter immediately before loading onto the MBPTrap HP column.
vi. Prior to loading the clarified lysate, equilibrate the MBPTrap HP column with 7 column volumes (CV) of binding buffer at a flow rate of 1 mL/min
vii. Load clarified lysate onto the column. The binding capacity of the 1 mL MBPTrap HP column is protein dependent but can bind approximately to 5 mg-7 mg of MBP-tagged protein. The flow rate should be decreased to 0.5 mL/min.
viii. Wash with 10 CV of binding buffer at a flow rate of 1 mL/min. If real-time A280 readings are possible, wash until no discernible absorbance at 280 nm is observed.
ix. Add 5 CV of elution buffer at a reduced flow rate of 0.5 mL/min, and assay elute fractions to determine fractions with the highest concentration of the protein of interest with a Bradford or BCA assay.
x. Regenerate the column with 3 CV distilled water followed by 3 CV of 0.5 M NaOH. Wash away the NaOH with 5 CV distilled water. The column is now ready to be used again.

## 4. Expected Results

The pFastBac-IRE1α-CD plasmid features are shown in **Figure 3** [20]. IRE1α-CD is fused to the C-terminus of an MBP protein under the polyhedrin promoter. A 6X His tag is also present at the N-terminal of the MBP protein. The MBP protein and IRE1α-CD coding sequences are separated by a Tobacco Etch Virus (TEV) protease site. The plasmid has an ampicillin (amp) and gentamicin (gent) selection marker for selection during the preparation of recombinant bacmid. The pFastBac backbone has a Tn7att transposition element that guides the insertion of the coding sequence into the bacmid. After transformation into *E. coli* DH10Bac, colonies were grown for 48 hrs. Colonies show up within 24 hrs but take an additional day to develop a blue color. Large white colonies were selected and re-streaked onto a Tet-Kan-Gent-IPTG-X-gal plate.

After this additional round of selection, a colony PCR was performed to make sure that IRE1α-CD was incorporated into the recombinant bacmid (**Figure 4**). Primers specific to an internal region in IRE1α-CD were used. As expected, a band was seen in all of the white colonies picked, indicating that the gene of interest was inserted in the bacmid. Bacmids were prepped and sent for DNA sequencing. Once the IRE1α-CD sequence was confirmed, glycerol stocks were made for future use (see **Supplemental file**).

The recombinant bacmid was extracted from *E. coli* DH10Bac cells and precipitated with 100% isopropanol to obtain pure bacmid samples. Sf21 cells were transfected and monitored for signs of infection for 5 days and multiple passages of viral titer. **Figure 5** shows the Sf21 cells changing on infection with P3 baculoviruses. The cells become larger in size, and the nuclei appear to occupy more of the cell. The Sf21 cells show more granularity as the infection progresses.

We checked the expression of IRE1α-CD after transfection by collecting cell lysate after P0 and P1 infections. **Figure 6** shows a GelCode Blue-stained SDS-PAGE (8%) of cell lysates after transfection and infection with baculoviruses to express IRE1α-CD. As evident in the figure, the IRE1α-CD protein is produced along with other contaminating insect cell proteins, also stained by GelCode Blue and therefore needs to be purified for downstream assays. The proportion of IRE1α-CD protein expressed in P1 infected cells was more than P0 infected cells, reflecting the higher concentration of baculoviral titer. The baculovirus stock was amplified to P3 to obtain high protein expressing infected Sf21 cells.

**Figure 6:**
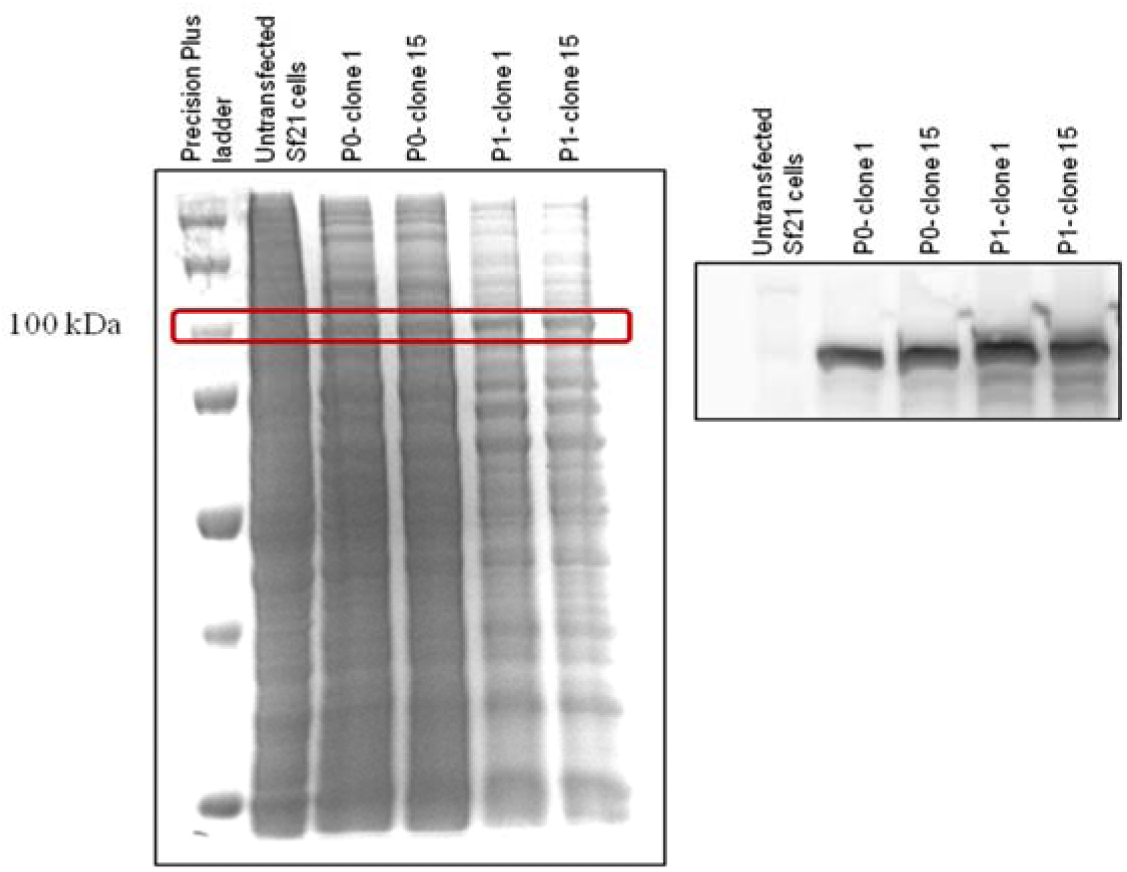
Expression of MBP-IRE1α-CD protein prior to purification. (Left) SDS-PAGE of total protein lysate for untransfected Sf21 cells, P0 and P1 infected clones 1 and 15. The 100 kDa band is marked and the putative 6X His-MBP-IRE1α-CD protein band is prominent in P1 infections. (Right) Western blot using an anti-IRE1 antibody

The P3 viral supernatant was collected, filtered with a 0.22 µm filter and stored at 4°C in the dark for subsequent infection. Sf21 cells were passaged the day before infection so that they were in log phase at a density of 3×10^5^ cells/mL. 150 µL of P3 supernatant was added for each mL of Sf21 cells. The time required for the expression of a protein depends on each protein of interest expressed.

**Figure 7** shows the western blot for the different time points throughout protein expression to determine the optimum time point for harvesting the cells. For IRE1α-CD, in P0 infected cells, maximum expression was observed at 96 hrs. Upon increasing infection titer by using P2 stocks, the overall level of protein expression increased. The optimal time point for harvesting was still after 96hrs of infection.

Once the viral titer and time for expression were optimized, the cell lysates were collected for purification of the protein. We used the MBP tag on the IRE1α-CD to aid in the purification. Filtered and clarified lysate was run through the MBPTrap column to obtain IRE1α-CD protein. The protein was of >85% purity based on densitometric analysis of the bands seen in an SDS-PAGE gel. Once IRE1α-CD is purified, it can be used for binding assays to determine binding partners for the protein [21]. **Figure 8** below shows the various fractions from the purification process outlined in the manuscript. The addition of an MBP-tag makes for a very specific purification. The yield of the expressed protein obtained after purification differs by the plasmids used, by baculoviral stocks and by the type of protein being expressed. The final volume of Sf21cells infected for scale-up can be adjusted based on the yield of protein required.

**Figure 7:**
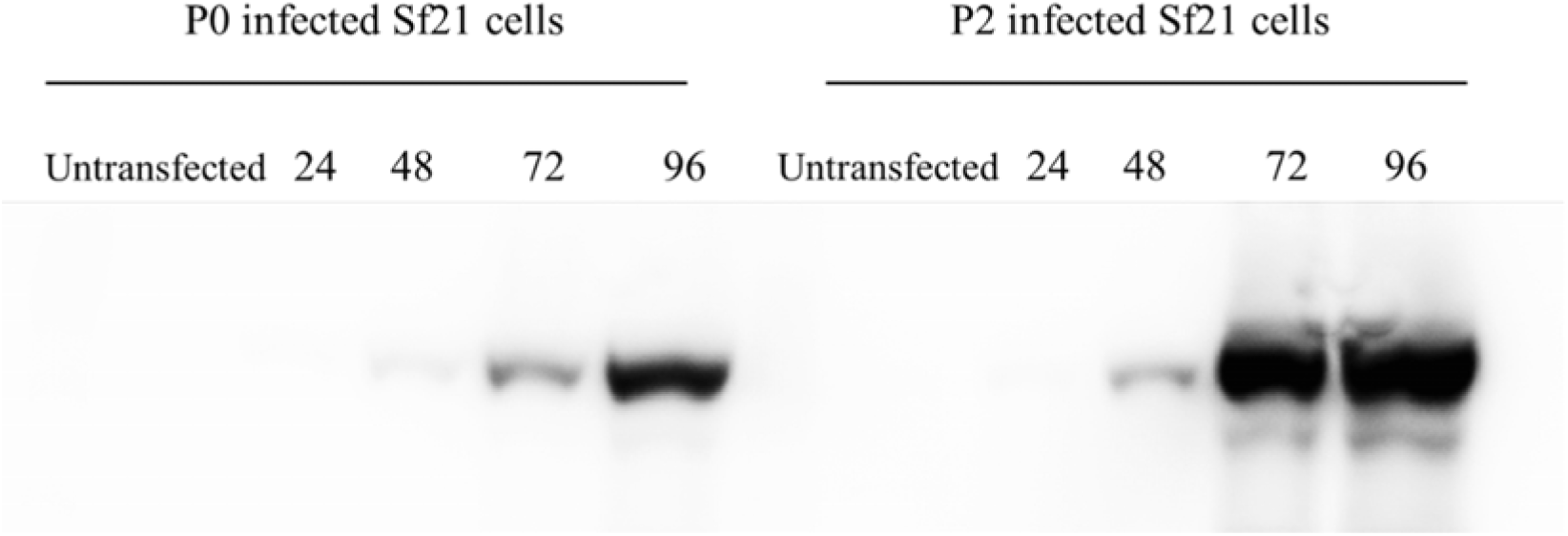
Expression of IRE1-CD protein. Western blot showing the time course of protein expression in P0 (left) and P2 (right) infected Sf21 cells using anti-IRE1α antibody

**Figure 8:**
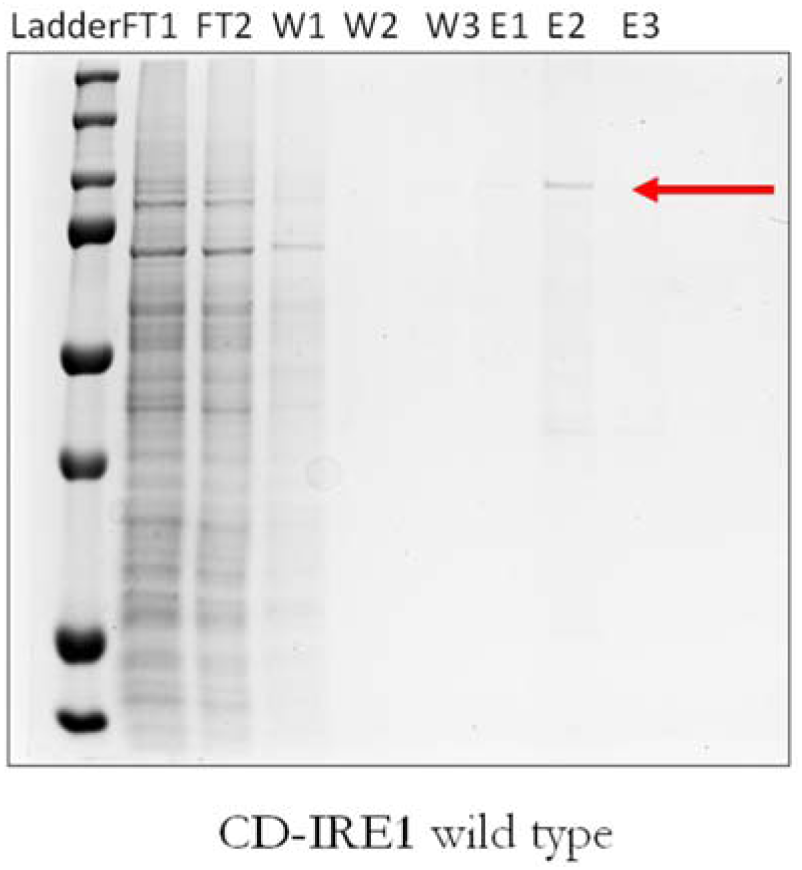
A GelCode Blue stained SDS-PAGE showing the various fractions from the purification procedure of IRE1-CD. FT-Flowthrough, W-Wash fractions, E-Elute fractions. The ladder is the Precision Plus blue ladder (Bio-Rad, cat# 1610373). The E2 fraction shows a pure protein band at ∼100kDa as expected

Sf21 insect cells combined with the pFastBac baculoviral system make a very robust, optimizable system for expression of eukaryotic proteins. MBP-IRE1α-CD was successfully expressed and purified from Sf21 cells. This protocol can be easily modified to express any mammalian protein in Sf21 insect cells by modifying the PCR primers shown in **Table 4** for the desired CDS. Long term expression of the protein can be achieved by making frozen stocks of baculovirus infected Sf21 cells. The MBP tag can be cleaved off using AcTEV protease if it interferes with downstream assays and the protein can be cleaned up using a Ni-NTA column. An AKTA-FPLC automated system can be used to aid in purification of higher volumes of cell lysate. If a secretion tag is used and the protein of interest is secreted in the media, a 5mL MBPTrap column can be used with the AKTA FPLC system to automate the purification process.

## 5. Summary

Further characterization of the expressed IRE1α protein is required to establish its structure, phosphorylation status and *in vitro* XBP1 splicing activity conclusively. The phosphorylation status of the protein can be identified by various methods. Feldman et al., used mass spectrometry techniques to identify the phosphorylated sites on IRE1α [22]. This technique in conjunction with a Phos-tag gel would lend more information on the ratio of phosphorylated to unphosphorylated species in the purified protein[23], [24]. Using the Phos-tag ligand developed by Fujifilm Wako-Chem, an SDS-PAGE gel can be used to separate phosphorylated proteins from unphosphorylated forms. The Phos-tag ligand binds to the phosphate group and slows down migration of the phosphorylated band. This leads to a separation of the phosphorylated and unphosphorylated species into separate bands. To confirm that the band pattern is because of distinct phosphorylation, purified protein samples can be treated with λ-phosphatase enzyme with activity towards phosphorylated serine, threonine and tyrosine residues. On treatment and subsequent application to a Phos-tag SDS-PAGE gel, the slower migrating band should disappear. *In vitro* XBP1 splicing can be assayed with a hairpin-RNA cleavage assay or by a fluorescence-based assay with a FRET-paired oligonucleotide [18].

If the protein structure needs to be determined, circular dichroism or small-angle X-ray scattering techniques can be used. Circular dichroism is a very useful tool to determine secondary structure as well as native folding of expressed or fusion proteins. Specifically, secondary structure can be determined using far-UV spectra and protein folding characteristics can be determined by monitoring spectra at different temperatures or in the presence of different denaturing agents [25].

## Author Contributions

Conceptualization, A.O. and C.C.; Methodology, A.O.; Validation, A.O., G.J. and C.C; Writing—original draft preparation, A.O.; Writing—review and editing, A.O., G.J, C.C.; Funding and Supervision, C.C.

## Funding

This research was funded by National Science Foundation, grant number CBET 1510895, CBET 1547518 and CBET 1802992.

## Acknowledgments

The authors would like to thank Dr. Xiangshu Jin, Michigan State University for providing the Sf21 cell line for this work.

## Conflicts of Interest

The authors declare no conflict of interest. The funders had no role in the design of the study; in the collection, analyses, or interpretation of data; in the writing of the manuscript, or in the decision to publish the results.

## Supplemental file

### I. Preparation of competent *E coli* DH10Bac cells

#### Materials needed

i. *E. coli* DH10Bac (streak from glycerol stock onto a Luria agar plate containing tetracycline/kanamycin/gentamycin, get isolated colonies and inoculate single colony) in LB broth containing tet/kan/gent grown at 37°C at 200 rpm.
ii. Ice cold centrifuge tubes, P10, P200, P1000 tips, microcentrifuge tubes and cryovials
iii. Refrigerated centrifuge
iv. Dimethyl sulfoxide (DMSO)
v. FSB buffer: (Sterile filter through a 0.22 µm filter and store at 4°C) 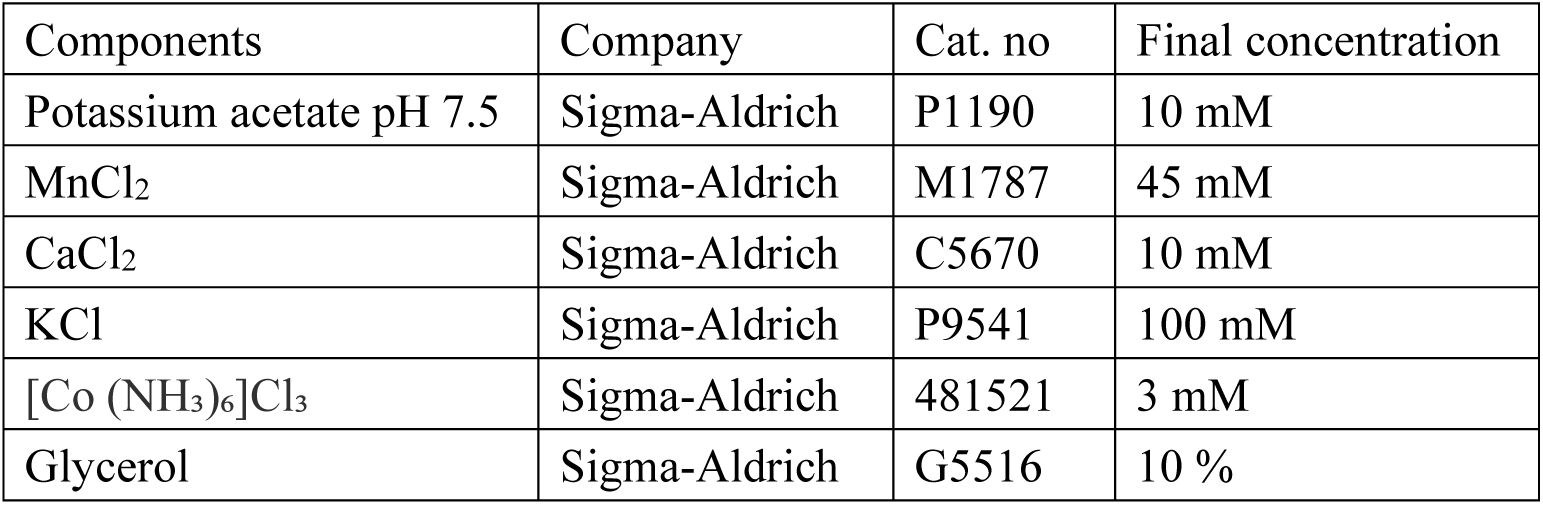

#### Protocol

i. Grow *E.coli* DH10Bac cells overnight until the OD is between 0.3-0.6. The LB media should contain kanamycin and tetracycline at the working concentrations listed in Table 8 of the main text. * *It is very important not to overgrow the culture*\*
ii. Cool the cells in wet ice for 10-20 minutes. \**From now on the cells should remain always ice cold*\*
iii. Pellet the cells by centrifugation, 10 min at 2500 rpm, 4°C.
iv. Resuspend the cells gently in 24 ml of ice-cold FSB.
v. Incubate on wet ice for 10-15 minutes.
vi. Pellet then cells again as above.
vii. Resuspend the cells in 8 ml of cold FSB.
viii. Add 280 uL DMSO.
ix. Incubate for 5 minutes on wet ice.
x. Add 280 uL DMSO.
xi. Incubate for further 5 minutes on wet ice.
xii. Aliquot the cells in 200-400 uL batches to sterile single-use cryovials.
xiii. Store in −80°C. \**Do not store in liquid nitrogen*\*

### II. Preparation of glycerol stocks for *E coli* DH10Bac cells with recombinant bacmid

#### Materials needed

i. Glycerol (Sigma-Aldrich, cat # G5516)
ii. Cryovials (Sigma-Aldrich, cat# 5000-0020)

#### Protocol

i. Prepare a 60% glycerol solution in water. Filter sterilize using a 0.22 µm filter or autoclave the solution at 121°C for 15 mins.
ii. Grow *E. coli* DH10Bac cells with recombinant bacmid in LB media for 14-16 hrs. The LB media should contain kanamycin, gentamicin and tetracycline at the working concentrations listed in **Table 8** of the main text.
iii. Add 750 µL of cells and 250 µL of the glycerol solution to a cryovial.
iv. Invert a few times to mix properly.
v. Store at −80°C.

### III. Preparation of recombinant bacmid from *E. coli* DH10Bac cells

#### Materials needed

i. PureLink HiPure Plasmid Miniprep kit (buffers R3, L7, N3) (ThermoFisher, cat# K210002)
ii. 100% isopropanol

#### Protocol

i. For recombinant bacmids, use 15–25 mL of an overnight *E. coli* DH10Bac with recombinant bacmid grown in Luria Broth.
ii. Pellet the cells at 5000 × g for 7 minutes to harvest the cells. Remove all medium and wash with PBS if necessary.
iii. Add 0.4 mL Resuspension Buffer (R3) with RNase A to the cell pellet in the tube and resuspend the cells. Gently shake the tube until the cell suspension is homogeneous.
iv. Add 0.4 mL Lysis Buffer (L7). Place the cap on the tube and ensure it is secure. Mix gently by inverting the capped tube until the lysate mixture is thoroughly homogenous. **Do not vortex**.
v. Incubate the lysate at room temperature for 5 minutes. **Do not exceed 5 minutes**.
vi. Add 0.4 mL Precipitation Buffer (N3) and mix immediately by inverting the tube until the mixture is thoroughly homogeneous. **Do not vortex**. Centrifuge the lysate at >12000g for 10 mins at RT. \**Don’t load onto the column provided with the kit. It decreases yield of bacmid and does not increase the quality of bacmid obtained*\*
vii. Take the supernatant gently without disturbing the pellet Add 0.63mL ice cold isopropanol to the supernatant. Mix well and incubate at RT for 15 mins. \**Alternatively, keep overnight at −20°C for precipitation*\*
viii. Centrifuge tube at >12000g for 30 mins
ix. Air dry the pellet for 10 mins, then resuspend the purified bacmid DNA in 60 uL water.
x. Solubilize bacmid DNA for 1 hr at 65°C or 4°C overnight. \**4°C overnight is much better than 65°C*\*
xi. Measure concentration of bacmid with Nanodrop.

